# Lymph node resident memory T cells retain effector capabilities by evading lung resident memory dysfunction

**DOI:** 10.64898/2026.02.03.703569

**Authors:** Taylor A Heim, Zachary J Rogers, Sarah Duquette, Molly Y Carney, Vidit Bhandarkar, Fiona Chatterjee, Teresa Bhandarkar, Jason M Schenkel, Scott Manalis, J Christopher Love, Stefani Spranger

## Abstract

Resident memory T cells (T_RM_) mediate localized immunity in barrier tissues while central memory T cells (T_CM_) recirculate through lymphoid organs to surveil for reinfection. Although T_RM_ are classically associated with peripheral non-lymphoid tissues, they have also been identified within lymph nodes (LN_RM_) where the mechanisms guiding their formation and functional differences remain poorly understood.

Here we used longitudinal antibody labeling to track the migratory history of memory T cells after influenza infection and demonstrate that CD69^+^CD103^+^ T cells are resident in the lymph node. LN_RM_ accumulate evenly throughout the lung-draining lymph node and are present within all analyzed LN compartments including, the sub capsular sinus, T cell zone and germinal centers. Epigenetic and transcriptional profiling reveal that LN_RM_ are uniquely poised for cytotoxicity whereas T_RM_ in the lung (Lung_RM_) resemble exhausted cells with elevated expression of inhibitory receptors and increased chromatin accessibility at the Pdcd1 locus. Regulatory network analysis of transcription factors, combined with target gene expression and chromatin accessibility, identified key regulons differentiating T_CM_, LN_RM_ and Lung_RM_ states. Upon antigen re-encounter, LN_RM_ are more proliferative, cytotoxic, and produce more IFNγ compared to Lung_RM_. Notably, we find that LN_RM_ represent the most prevalent subset of memory T cells in human thoracic lymph nodes. These findings highlight functional heterogeneity in T_RM_ and establish LN_RM_ as a distinct and durable memory T cell population bridging features of circulating and tissue-resident cells.

## Introduction

Memory T cells provide long lasting protection against previously encountered pathogens. They can be broadly divided into circulating populations, which patrol lymphoid tissues, and tissue-resident memory T cells (T_RM_), which persist at sites of prior infection. T_RM_ are well described in barrier tissues such as the lung, skin, gut, and brain, where they mediate rapid local responses^1–5^. In contrast, lymph nodes are traditionally considered domains of recirculating central memory T cells (T_CM_), with little contribution from resident populations. However, recent studies have identified bona fide T_RM_ in secondary lymphoid organs, including the spleen, skin-draining LN, and lung-draining LN (mediastinal LN; medLN), raising new questions about their function and origin^6–9^.

In murine models of vaccinia or influenza infection, LN_RM_ formation is dependent upon the migration of activated T cells from the virally infected non-lymphoid tissue into the draining lymph node^6,8^. Whether LN_RM_ share functional properties with classical barrier T_RM_ or represent a functionally distinct lineage remains unknown. Dissecting functional differences between LN_RM_ and T_CM_ is challenging, as lymph nodes are heavily populated by rapidly circulating memory T cells. Current markers such as CD69 and CD103 only correlate with and do not prove residency^1^. Previously, CD69^+^ CD8^+^ T cells were presumed to be responding to residual depots of antigen in the medLN after influenza infection^10^, confounding the distinction between activated and resident cells. Indeed, bona fide migration studies such as parabiosis, are required to determine the circulatory or resident status of a T cell. However, while parabiosis is the current gold standard for assessing tissue residency, the technique is technically demanding, inflammatory, and low throughput. Thus, the existence, regulation, and function of LN_RM_ populations remain understudied and less well characterized relative to circulating memory T cell populations.

To address these gaps, we used longitudinal intravascular labeling to track the dynamics of influenza-specific CD8^+^ T cells in blood and lymphoid tissues. We identified a population of CD69^+^CD103^+^memory T cells that stably resides in the medLN for months after infection. These LN_RM_ are highly conserved across multiple influenza strains, TCR specificities, and precursor types, suggesting that LN_RM_ formation is a generalizable feature of antiviral memory. Spatial mapping revealed that LN_RM_ localize broadly across distinct lymph node compartments, including the subcapsular sinus, B cell follicles, germinal centers, and high endothelial venules.

Using integrated single-cell RNA and ATAC sequencing, we show that LN_RM_ are transcriptionally and epigenetically distinct from both Lung_RM_ and T_CM_. LN_RM_ lack expression of recirculation machinery and instead constitutively express cytotoxic proteins, including granzymes. Functionally, LN_RM_ display superior proliferative and effector potential compared to Lung_RM_. These findings reveal the existence of a stable, highly functional population of resident memory T cells in the lymph node, with implications for vaccine design in the respiratory tract and immunotherapies.

## Results

### Longitudinal antibody labeling distinguishes circulating and resident memory T cells in the mediastinal lymph node

To study memory T cell distribution following viral infection, we transferred naïve CD90.1^+^ OT-I T cells into naïve C57BL6 mice and infected mice with intranasal influenza virus expressing SIINFEKL(PR8-OVA). At 21 days post infection, we detected OT-I T cells within blood, secondary lymphoid organs and the lung. As expected^11–14^, most cells in the lung expressed CD69 and CD103, canonical markers of tissue residency while T cells in the blood and most secondary lymphoid organs were CD69^-^CD103^-^ (**Fig 1A, Fig S1A**). Interestingly, 66% of T cells in the lung-draining mediastinal lymph node (medLN) expressed CD69 and CD103, comparable to the proportion of CD69^+^CD103^+^ T cells found in the lung (**Fig 1A**).

**Fig 1.**
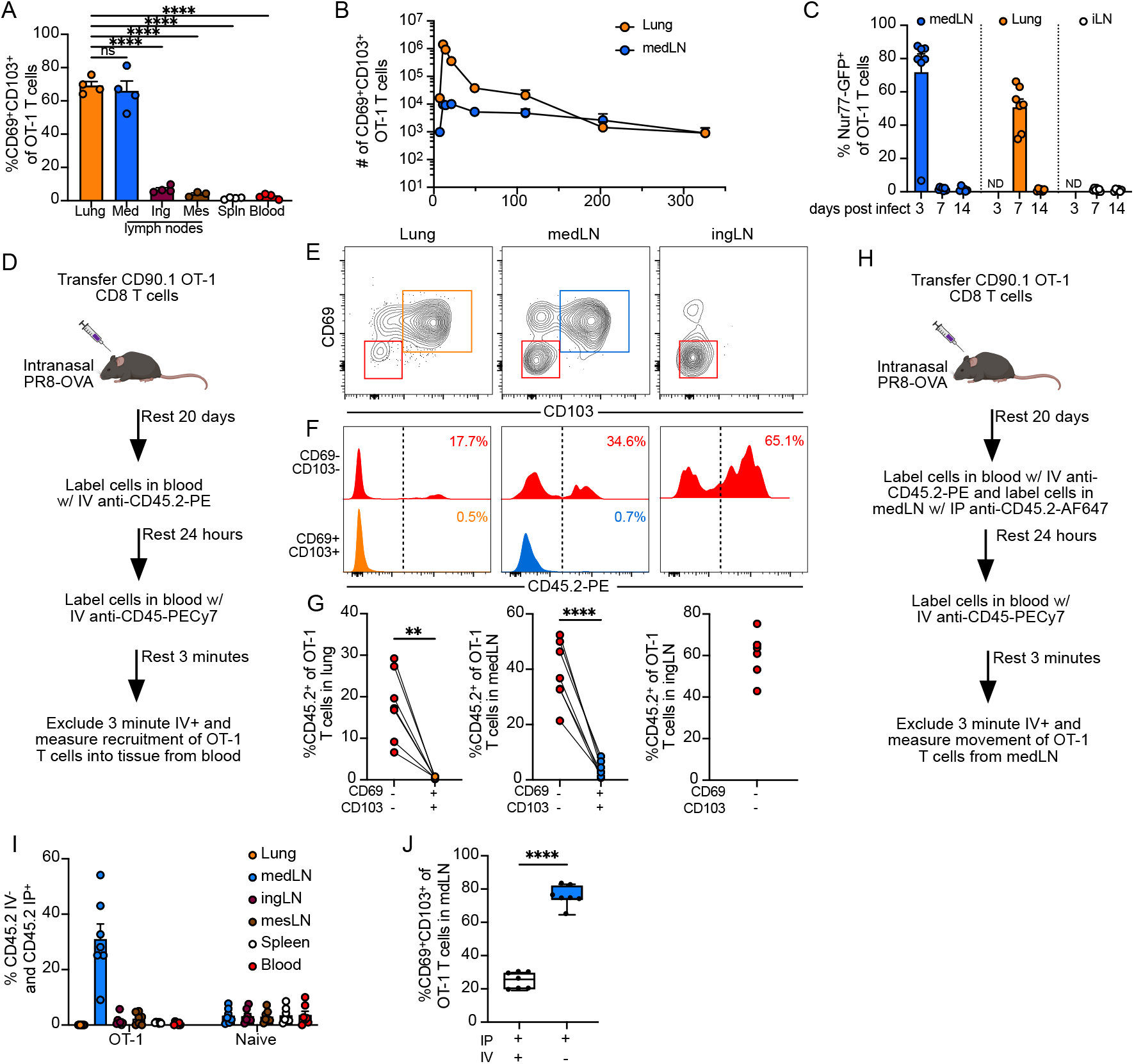
Longitudinal antibody labeling distinguishes circulating and resident memory T cells in the mediastinal lymph node. (A) Naive CD90.1+ OT-I T cells were transferred into naive B6 mice before intranasal infection with influenza expressing SIINFEKL (PR8-OVA). Expression of CD69 and CD103 was measured 21 days post infection on extravascular cells. (B) Quantification of CD69^+^CD103^+^ OT-I T cells. (C) Naïve CD90.1^+^ OT-I Nur77-GFP T cells were transferred into mice one day before intranasal PR8-OVA infection. Nur77-GFP expression was evaluated 3-14 days later by extravascular OT-I T cells. (D)Experimental setup for (E-G). Flow cytometry plot (E) of CD90.1^+^ OT-I T cells isolated from indicated locations. (F-G) Longitudinal labeling of cells and quantification. (H) Experimental setup for (I-J). (I) Labeling after intravenous and intraperitoneal injection of OT-I T cells or naïve endogenous T cells. (J) Expression of CD69 and CD103. Naïve= CD8^+^CD90.1^-^CD44^low^. Each point represents a mouse and N=2-5 mice per experiment. Data are representative (A) or are pooled from at least two experiments (n=2–4/group), except for day 113 in (B), which reflect a single experiment.

Resident memory T cells in the lower respiratory tract are poorly maintained over time relative to other anatomical locations^12,13,15^. In prior reports, LN_RM_ in the medLN following influenza infection were durably maintained up to 120 days^8^ but in separate experiments using LCMV, LN_RM_ appeared to be entirely lost from the medLN between 60 and 400+ days post infection^16^. Thus, we interrogated the persistence of T_RM_ across both sites. LN_RM_ contracted only 3.8-fold between day 21 and 203 while Lung_RM_ contracted 249.0-fold over the same time frame, confirming previous reports on the short-lived nature of Lung_RM_ and durability of LN_RM_ (**Fig S1B,C**). We extended our analyses out to 326 days post infection and observed continual attrition of Lung_RM_ while LN_RM_were numerically stable (**Fig 1B**). CD69^+^CD103^+^ OT-I T cells were almost 100-fold more prevalent in the medLN after accounting for differences in mass (**Fig S1D**) after 100 days post infection, highlighting the abundance of T_RM_ in the medLN relative to the lung.

The presence of CD69^+^ T cells in the medLN after viral clearance has been attributed to persistent antigen depots^10,17^, and we postulated that the maintenance of LN_RM_ might be due to sustained antigen recognition in the medLN. To address this, we utilized Nur77-GFP transgenic mice in which T cells express GFP upon TCR engagement with cognate peptide:MHC^18^ and crossed them to OT-I mice. As expected, transferred Nur77-GFP OT-I T cells in the medLN expressed GFP at day 3 post-infection, consistent with robust TCR signaling during the priming phase (**Fig 1C, Fig S1E**). At day 7, transferred T cells in the lung were ∼50% GFP^+^ but we were surprised to see minimal GFP expression in the medLN. The low percentage of GFP^+^ cells in the medLN indicated that there was either no antigen in the lymph node or that the Nur77-GFP OT-I T cells were not encountering antigen presenting cells. Signs of active antigen recognition in both the lung and lymph nodes were absent by day 14 (**Fig 1C)** indicating that LN_RM_ did not encounter deposits of antigen at either organ site.

While resident memory T cells have been reported in secondary lymphoid organs^6– 9,19^, most memory CD8^+^ T cells in lymph nodes are presumed to be circulating memory which led us to investigate whether CD69^+^CD103^+^ T cells in the medLN were indeed resident. To distinguish between OT-I T cells in the medLN that were recently recruited from blood or resident memory T cells, we fluorescently labeled all leukocytes in the blood with anti-CD45.2-PE via intravenous injection^20,21^. 24 hours later, we quantified the number of fluorescently labeled cells that had moved into tissues from the blood vasculature. To discriminate extravascular T cells from intravenous T cells, we injected anti-CD45, labeled with PECy7, 3 minutes before euthanizing mice^22^ (**Fig 1D**). The CD69^+^CD103^+^ T cells in the lung and medLN were almost exclusively negative for anti-CD45.2-PE, suggesting that these T cells were resident for at least 24 hours (**Fig 1E-G**). As a reference point, we assessed OT-I T cells in the inguinal lymph node, which lacked CD69 and CD103 marker expression. On average 61% of OT-I T cells in inguinal lymph nodes were labeled with CD45.2-PE, indicating recent recruitment from blood (**Fig 1E-G**).

The prior experiments allowed us to conclude that CD69^+^CD103^+^ T cells found in the medLN were not recently recruited from blood but did not necessarily prove that these T cells were resident in the medLN. We next used a similar longitudinal labeling technique^20,21^ to label cells in the medLN via intraperitoneal (IP) injection of anti-CD45.2-AF647. A simultaneous injection of anti-CD45.2-PE intravenously allowed us to distinguish between T cells that were in the medLN or blood at the time of injection (**Fig 1H**). 24 hours later, 31% of OT-I T cells in the medLN were positive for the IP label while being negative for the IV label (IP^+^IV^-^; **Fig 1I, Fig S1F**). This observation indicates that these OT-I T cells had been in the medLN at the time of injection and remained resident for at least 24 hours (**Fig 1I**). We also assessed naïve CD8^+^ T cells across lymph nodes and observed equal labeling (IP^+^IV^-^) across all lymph nodes, including the medLN, suggesting that naïve T cells were continuously circulating in and out of the medLN (**Fig 1I**). In addition, IP^+^IV^-^ OT-I T cells in the medLN were mostly CD69^+^CD103^+^ which further supports our conclusion that the IP^+^IV^-^ T cells in the medLN are bona fide resident memory T cells (**Fig 1J**). This finding is in agreement with previous parabiosis experiments in which Ly6C^-^CD103^+^ memory T cells in the medLN failed to equilibrate between host and partner^8^. Thus, longitudinal antibody labeling represents a valuable, non-invasive tool to infer the migratory status of a T cell and demonstrates the residency of CD69^+^CD103^+^ T cells in the medLN after influenza infection.

### LN_RM_ differentiation is conserved across influenza strains, antigen specificities and precursor populations

The magnitude of T_RM_ in the medLN led us to question whether this observation was restricted to the PR8 strain of influenza or a consequence of the OT-I TCR transgenic model. We transferred naïve OT-I T cells into mice and infected intranasally with either PR8-OVA (H1N1) or X31-OVA (H3N2) (**Fig 2A**). Although X31 causes a milder infection compared to PR8^23^, both infections drove formation of CD69^+^CD103^+^ OT-I T cells in the medLN 26-27 days post infection. While OT-I T cells in the medLN after PR8-OVA infection were more likely to be CD69^+^CD103^+^ on a per cell basis (**Fig 2B, Fig S2A**), the total number of CD69^+^CD103^+^ OT-I T cells was strikingly similar following infection with either strain (**Fig 2C**).

**Figure 2.**
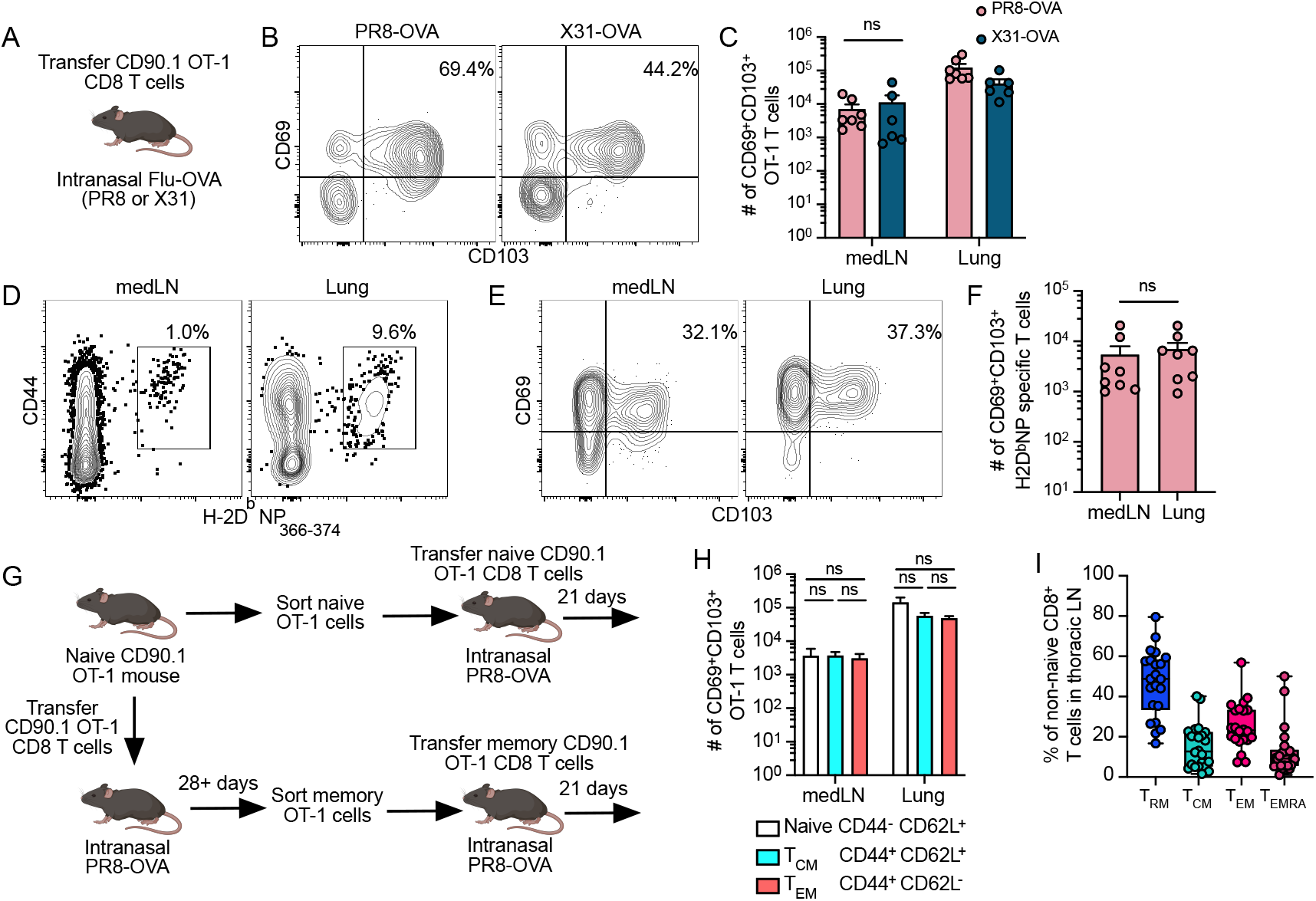
LN_RM_ differentiation is conserved across influenza strains, antigen specificities and precursor populations. (A)Experimental setup for B-F.(B-C) Representative flow plots of OT-I T cells in the medLN and quantification at 26-27 days post infection. Gated on CD90.1^+^, CD8^+^, CD44^hi^, live. n=6 or 7 combined from two experiments. (D) Representative flow plots of extravascular CD8^+^ T cells in lungs and medLN(D) 26-27 days after PR8-OVA infection. (E) Same as D but gated on tetramer^+^ cells. n=8 combined from two experiments. (F) Quantification of E. (G-H) Experimental setup where naïve or memory OT-I T cells were flow sorted and transferred into naïve mice before intranasal PR8-OVA infection. (I) Analysis of scRNA+CITEseq of 24 human thoracic lymph nodes adapted from Wells et al. 2025. See methods for subset definitions.

To address potential caveats of adoptively transferred TCR transgenic OT-I T cells, we also assessed endogenous CD8^+^ T cells specific for viral nucleoprotein (NP) (**Fig 2D**) following PR8-OVA or X31-OVA infection using a H-2D^b^:NP_366-374_ tetramer. In both the lung and the medLN about a third of all NP-reactive T cells were CD69^+^CD103^+^ (**Fig 2E, Fig S2B-C**). Despite the differences in organ size, we found comparable numbers of CD69^+^CD103^+^ T cells within the medLN and the lung (**Fig 2F**). Together these findings demonstrate robust induction of LN_RM_ across multiple viral strains and antigens.

Prior reports have demonstrated a variation in the ability of naïve or memory T cells to differentiate to resident memory T cells. In the small intestine following LCMV infection, naïve T cells seed more T_RM_ in the small intestine compared to T_EM_ or T_CM_^24^ while in human foreskin graft models T^CM^ are the dominant cell population seeding T_RM_^25^. Thus, we next investigated the extent to which different precursor populations could form LNRM. We transferred naïve OT-I T cells into CD90.2 host mice and infected mice with PR8-OVA. After at least 28 days post infection, we isolated T_CM_ (CD44^+^CD62L^+^) and T_EM_ (CD44^+^CD62L^-^) OT-I T cells from the spleen and transferred 20,000 cells intravenously to naïve CD90.2 host mice. To compare secondary memory responses to primary memory responses we also transferred naïve OT-I T cells (CD44^-^CD62L^+^) into CD90.2 host mice (**Fig 2G**). Interestingly all transferred T cell populations seeded similar numbers of CD69^+^CD103^+^ cells in both the lung and medLN, though naïve T cells transfer resulted in slightly more CD69^+^CD103^+^ cells in the lung (**Fig 2H**). To extend these findings to humans, we re-analyzed a single-cell RNA sequencing atlas of human lymphocytes in multiple organs, including lung-draining thoracic lymph nodes^26^. In thoracic lymph nodes, LN_RM_ comprised 48% of all non-naïve conventional CD8^+^ T cells (**Fig2I**). LN_RM_ were the most prevalent memory T cell subset—exceeding the frequency of central memory cells in all analyzed lymph nodes (thoracic, inguinal and mesenteric) (**Fig S2D-F**). These data emphasize lymph nodes as niches for T_RM_ and demonstrate robust plasticity in memory T cell differentiation towards a LN_RM_ program.

### Influenza-reactive LN_RM_are evenly distributed across the subcapsular sinus, T cell zone and B cell follicle

Previous descriptions of the spatial distribution of LN_RM_ after LCMV and vaccinia infections have demonstrated that these cells are not solely restricted to the T cell zone but rather are distributed throughout different lymph node compartments. After LCMV infection 70% of the CD8^+^CD69^+^ viral specific T cells were in the lymph node sinus^7^, while after vaccinia infection, CD103^+^ LN_RM_ preferentially localized near LYVE1^+^ lymphatic endothelial cells within skin draining lymph nodes^6^. To understand and quantify the spatial distribution of LN_RM_ after influenza, we used immunofluorescent imaging of Thy1.1^+^ OT-I T cells in medLNs following PR8-OVA infection (**Fig 3A**). By immunofluorescent imaging, over 80% of OT-I T cells in the medLN expressed CD103 (**Fig 3B**) which is a larger fraction than observed by flow cytometry, potentially indicating that CD103^-^ cells are preferentially isolated when preparing single cell suspensions for flow cytometry as has been proposed for non-lymphoid tissues^27^. Most OT-I T cells in the medLN expressed CD103, regardless of their location within the lymph node (**Fig 3C**), further supporting prior findings that LN_RM_ are broadly distributed across diverse lymph node compartments. Assessing the fraction of CD103^+^OT-I T cells within each compartment, we identified that 62% localized to the T cell zone and interfollicular areas while 18% were in the B cell zone, 13% were near lymphatics and 7% were in the subcapsular sinus (**Fig 3D**). However, these different compartments vary vastly in size. To account for differences in area, we quantified CD103^+^cell density across the medLN and found that LN_RM_ are distributed evenly across all four analyzed compartments (**Fig 3E**). Interestingly, CD103^+^OT-I T cells were also detected within active germinal centers, raising the possibility of interactions between LN_RM_ and B cells (**Fig 3F**). Pathogens can access lymph nodes via both the subcapsular sinus and high endothelial venules (HEVs). We examined these sites and found CD103^+^OT-I T cells positioned in the subcapsular sinus and adjacent to HEVs (identified via peripheral node addressin; PNAD), suggesting a potential role for LN_RM_ in immunosurveillance at key points of pathogen entry (**Fig 3F**). These findings reveal a surprisingly homogeneous distribution of LN_RM_ within the medLN following influenza infection, potentially reflecting an ability of LN_RM_ T cells to rapidly coordinate secondary immune responses across diverse immune cell types.

**Figure 3.**
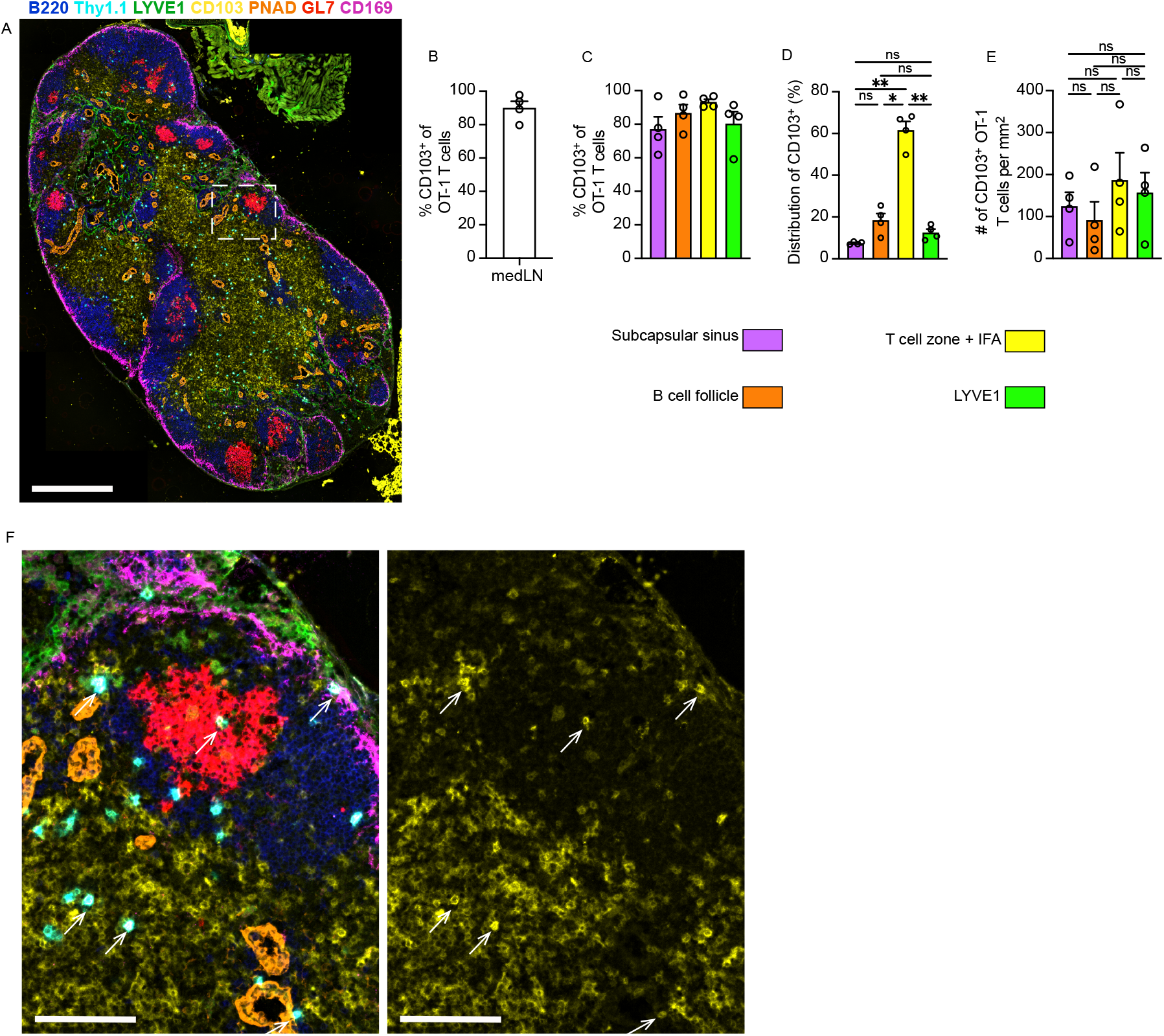
Influenza-reactive LN_RM_ are evenly distributed across the subcapsular sinus, T cell zone and B cell follicle. (A) Representative immunofluorescent imaging of medLN 45 days post PR8-OVA intranasal infection. (B-E)Quantification of CD103 expression in the entire medLN (B) or within select compartments (C-E). (F) Representative imaging of CD103^+^ OT-I T cells. Scale bar is 500μm (A) or 100μm (F). Data are from 4 mice from 38-45 days post infection.

### LN_RM_ are transcriptionally and epigenetically distinct from T_CM_ and Lung_RM_

Given that LN_RM_ are positioned in the lymph node but bear a resident memory phenotype, we questioned if these cells were more functionally related to Lung_RM_ or T_CM_. To understand differences between T_CM_, LN_RM_, and Lung_RM_, we isolated memory OT-I T cells from medLNs and lungs 46 days post influenza infection for single-nucleus RNA+ATAC sequencing (**Fig 4A**). This approach allowed us to compare transcriptional and epigenetic programs of T cells localized to the lung and medLN (**Fig 4A**) with the expectation that T_RM_ in both tissues would cluster closely together while exhibiting subtle, tissue-specific adaptations^28,29^.

**Figure 4.**
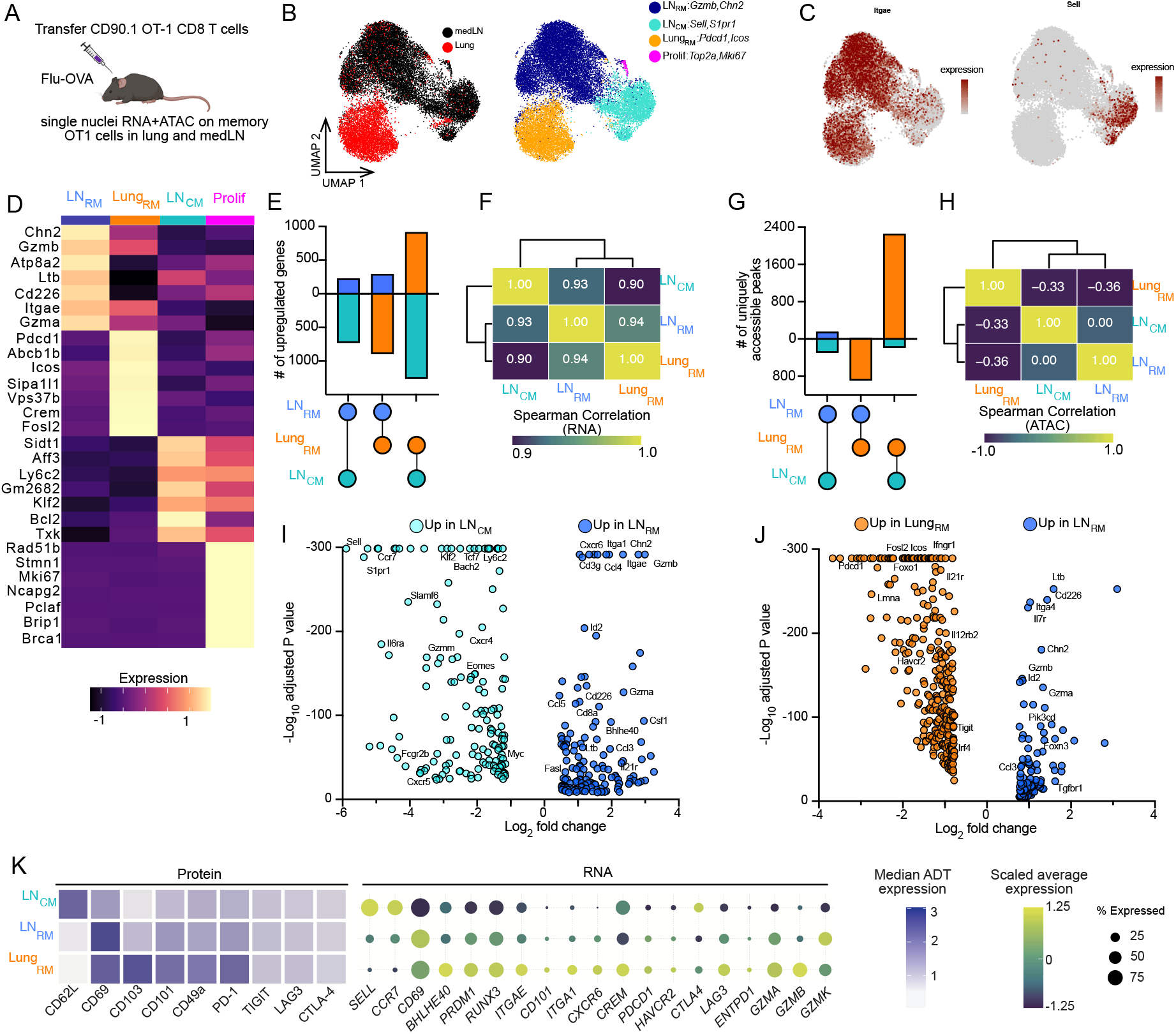
LN_RM_ are transcriptionally and epigenetically distinct from T_CM_and Lung_RM_. (A) Memory OT-I T cells were flow sorted (CD8^+^CD44^+^LiveDead^-^CD45IV^-^CD90.1^+^) from lung and medLN 46 days post infection. Nuclei were isolated for paired single nuclei ATAC+RNA sequencing. (B) UMAP of 14,866 nuclei denoting tissue of origin (left) or cluster (right). (C) Gene expression of *Sell (*CD62L) or *Itgae* (CD103). (D) Heatmap of top differentially expressed genes from each cluster. Number of differentially expressed genes(E) and Spearman analysis of similarity based on RNA expression(F). Quantification of chromatin peaks (G) and Spearman analysis of similarity based on ATAC(H). (I-J) Volcano plots of differentially expressed genes. (K) Protein and RNA expression on human CD8^+^ T cells from thoracic lymph nodes or lung from Wells et al. 2025. See methods for subset definitions.

Analysis of transcriptional data identified 3 large clusters and one smaller cluster expressing proliferation associated transcripts (**Fig 4B-D**). Cells derived from the medLN segregated into two distinct clusters with one expressing central memory-associated transcripts (LN_CM_; *S1pr1* and *Sell*) and a second cluster expressing transcripts associated with resident memory^3,30–32^ (LN_RM_; *Itgae* and *Cxcr6*) (**Fig 4B-D, Fig S3A-B**). Lung-derived T cells were mainly found in a single cluster (Lung_RM_) that expressed high levels of *Tox, Itgae* and inhibitory receptors *Pdcd1* and *Ctla4*. Both T_RM_ clusters expressed low amounts of transcripts needed for T cell circulation such as *S1pr1, Sell*, and *Ccr7* (**Fig 4B-D**). The largest difference in transcriptional programs was between Lung_RM_ and LN_CM_ with 2183 differentially expressed genes (1266 upregulated by Lung_RM_ vs 917 upregulated by LN_CM_) (**Fig 4E**). Spearman correlation analysis of RNA sequencing revealed that the transcriptional profile of LN_RM_ occupies an intermediate position between the Lung_RM_ and LN_CM_ (**Fig 4F**). Similarly, the greatest disparity in chromatin structure was found between Lung_RM_ and LN_CM_ with a total of 2444 differentially accessible peaks (2256 in Lung_RM_ vs 188 by LN_CM_) (**Fig 4G**). Spearman correlation analysis of ATAC data revealed that the epigenetic profile of LN_RM_ more closely aligns with LN_CM_ (**Fig 4H**). The pronounced disparity in gene expression and chromatin accessibility between LN_CM_ and Lung_RM_ highlight how T cell states are shaped by core transcriptional programs and tissue-specific adaptations.

When compared to LN_CM_, T_RM_ in the medLN increased gene expression of integrins (*Itgae* and *Itga1*), chemokines (*Ccl3, Ccl4*, and *Ccl5*), and genes supporting TCR signaling (*Cd8a* and *Cd3g*) (**Fig 4I**). Conversely, LN_CM_ expressed high levels of genes promoting T cell circulation (*Sell, S1pr1*, and *Ccr7*) as well as transcription factors *Klf2* and *Tcf7* (**Fig 4I, Fig S3A**). Comparing T_RM_ in the lung to the LN counterparts we observed that Lung_RM_ were defined by high expression of inhibitory receptors (*Pdcd1, Havcr2*, and *Tigit*) (**Fig 4J, Fig S3A**) potentially indicating reduced cytotoxic capability. LN_RM_ expressed the highest levels of *Gzmb* and *Gzma* relative to both Lung_RM_ and T_CM_, suggesting a heightened killing capacity by LN_RM_ (**Fig 4D,I,J Fig S3A**).

To determine whether these differences represent evolutionarily conserved features of T cell memory, we interrogated a publicly available multimodal dataset (scRNA-seq and CITE-seq) of CD8^+^ T cells in human lungs and thoracic lymph nodes^26^. Consistent with our murine findings, human Lung_RM_ expressed the highest levels of PD-1 protein and transcripts for inhibitory receptors such as *PDCD1, HAVCR2*, and *LAG3* (**Fig 4K**). We observed slight differences in granzyme expression patterns by human T cells. Human Lung_RM_ expressed the highest transcriptional levels of *GZMA* and *GZMB*, while LN_RM_ produced the highest GZMK, yet both T_RM_ populations expressed higher levels of granzymes than LN_CM_ (**Fig 4K**). These findings further highlight that T_RM_ appear better poised for cell killing due to high basal granzyme expression, while also identifying high inhibitory receptor expression as a distinct feature of Lung_RM_.

### LN_RM_ are more durable than Lung_RM_, do not rely on persistent antigen encounter and lack state-defining transcription factor activity

Because *Gzmb* was among the top differentially expressed genes by LN_RM_, we examined the epigenetic regulation of cytotoxicity (**Fig 4D**). While the *Gzmb* locus was least accessible in LN_CM_, we observed minimal differences in accessibility between Lung_RM_ and LN_RM_ (**Fig 5A**). Despite this epigenetic similarity, LN_RM_ expressed the highest levels of Granzyme B protein, mirroring their high transcript levels (**Fig 5A-B**). This discordance suggests that LN_RM_ are uniquely poised for granzyme-mediated cytotoxicity through post-transcriptional regulation, distinct from the regulation in Lung_RM_.

**Figure 5.**
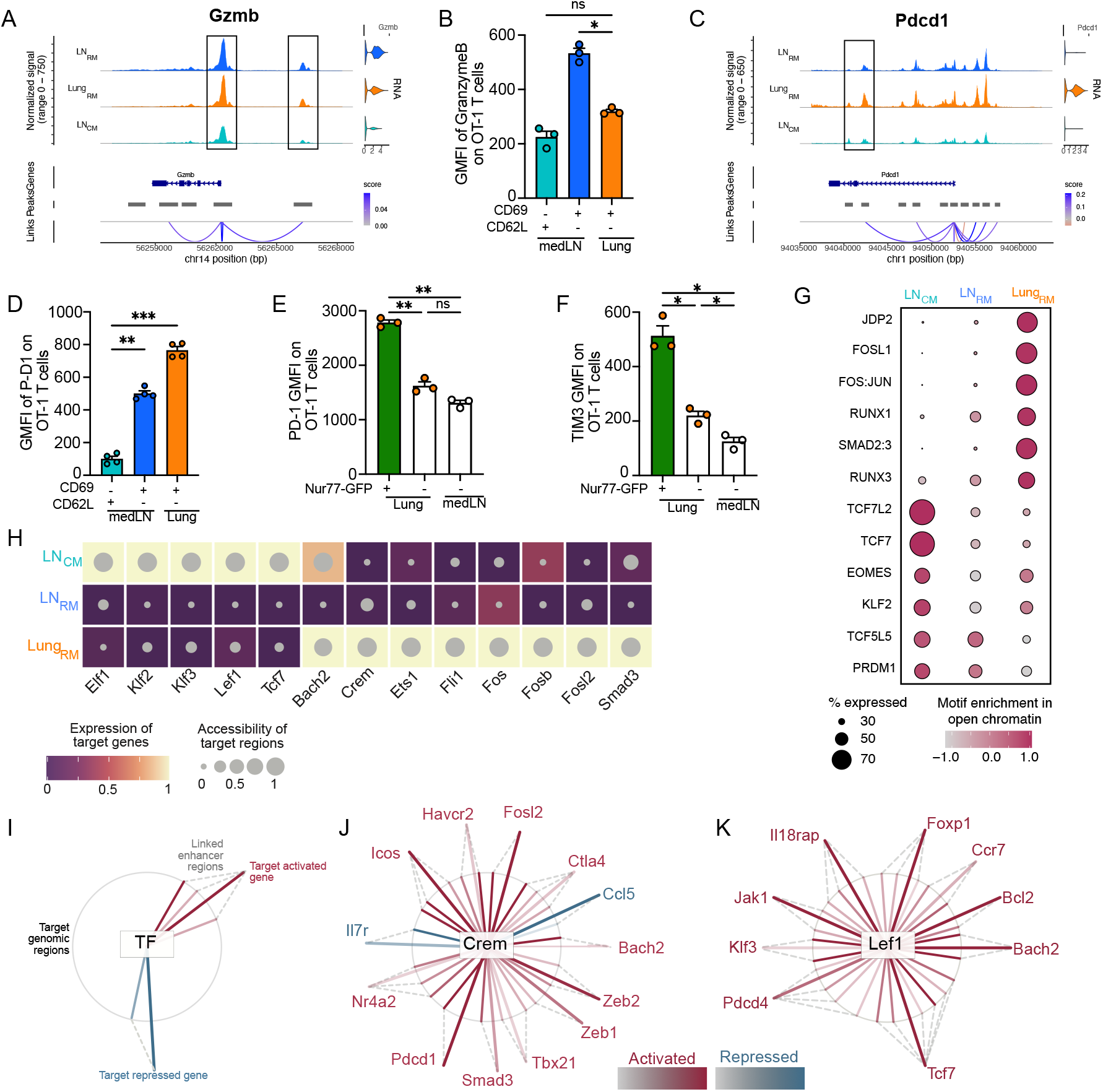
LN_RM_ are more durable than Lung_RM_, do not rely on persistent antigen encounter and lack state defining transcription factor activity. (A)Chromatin accessibility with gene expression (right) and linked chromatin peaks (bottom) at the *Gzmb* locus. (B) Protein expression of GranzymeB as measured by flow cytometry. (C-D) Same as (A-B) but for the *Pdcd1* locus. Protein expression on OT-I T cells is 49 days post infection where each dot represents one mouse. (J and L) Data are from one experiment and representative of at least two experiments with N=3. (E-F) Surface expression of PD-1 and TIM3 at 7 days post infection. (G) Transcription factor motif footprint analysis from Fig 4 dataset. (H) Inferred eRegulons identified by SCENIC+ across memory T cells after influenza infection. (I-K) Legend (left) for network plots illustrating inferred transcription factor relationships to target genes. Transcription factor and gene relationship is denoted by color of line represents activation (red) or repression (blue).

We next examined PD-1, a critical negative regulator of T cell effector function. Given the high *Pdcd1* expression observed in Lung_RM_, we analyzed chromatin accessibility at the Pdcd1 locus. Lung_RM_ displayed the greatest accessibility at the Pdcd1 locus, alongside the highest gene and surface protein expression. In contrast, LN_RM_ and LN_CM_ showed minimal Pdcd1 accessibility and expression (**Fig 5C-D**). These data define LN_RM_ as a distinct T cell state, possessing the cytotoxic potential of effector-like cells without the inhibitory profile of Lung_RM_. To determine if the elevated PD-1 in Lung_RM_ was driven by antigen recognition in the lung, we utilized OT-I Nur77-GFP reporter mice. In the lung at day 7 post infection, GFP^+^ (antigen-experiencing) OT-I T cells expressed the highest levels of PD-1 and TIM3 (**Fig 5E-F, Fig S4A**). This correlation suggests that secondary antigen encounters in the infected lung environment drive sustained inhibitory receptor expression that persists into the memory phase.

To understand transcriptional drivers of these states, we analyzed open chromatin regions for transcription factor motif enrichment. Lung_RM_ had enhanced chromatin accessibility at sites bound by RUNX proteins, which have well established roles in promoting T_RM_ differentiation^31^. Interestingly, Lung_RM_ were also enriched for AP-1 associated transcription factors (JDP2, FOSL1, FOS: JUN) downstream of cytokine or TCR induced activation (**Fig 5G**). In contrast, LN_CM_ showed high degrees of enrichment for TCF/LEF family of transcription factors (TCF7 and TCF7L2) (**Fig 5G**). LN_RM_ lacked unique motif enrichment and were characterized by a hybrid chromatin architecture sharing RUNX motifs with Lung_RM_ and TCF motifs with LN_CM_ (**Fig 5G**).

Transcription factor foot printing from ATAC seq is useful in understanding where transcription factors can bind but does not reveal downstream target genes and regulatory networks. To overcome this limitation, we used SCENIC+^33^ to extrapolate enhancer driven gene regulatory networks or eRegulons (TF+target genes) by connecting TF binding sites with observed gene expression. eRegulons are assembled by identifying chromatin accessible enhancer regions and identifying genes, including TFs that tend to be co-expressed together. For each co-expression module, the prevalence of TF motifs within enhancer regions is analyzed and used to score TF activity in conjunction with gene module activity. UMAP visualization of cells embedded by eRegulon activity clustered cells by cell type corresponding to LN_CM_, LN_RM_, and Lung_RM_ (**Fig S4B**). **Fig. 5H** integrates transcription factor target gene expression and chromatin accessibility at TF-bound enhancer regions. LN_CM_ were governed by KLF2, KLF3, TCF7 and LEF1 networks. Lung_RM_ were driven by AP-1 (FOS, FOSB, FOSL2), CREM and SMAD3 activity. Clustering of cells based upon gene regulatory networks activity, resolved three populations corresponding to LN_CM_, LN_RM_, and Lung_RM_ (**Fig 5H, S4C**). LN_RM_ did not score highly for any unique eRegulons, suggesting that LN_RM_ lack state defining transcription factor networks.

We next analyzed the inferred target genes for the identified TFs (**Table S1**) and visualized them using network plots (**Fig 5I-K**). The CREM eRegulon was highly active in Lung_RM_ and was inferred to drive expression of many inhibitory receptor genes including *Pdcd1, Havcr2*, and *Ctla4* while repressing *Il7r* (**Fig 5J**). Many of the identified TFs regulated expression of other highly expressed TFs, demonstrating that T cell states are not defined by a single TF but are products of complex gene regulatory networks. For instance, the LEF1 target genes included Tcf7 and Klf3 which were both highly active in LN_CM_ (**Fig 5K**). Consistent with prior analysis of T cell gene regulatory networks^32^, we found that KLF2 drove *S1pr1, Sell*, and *Ccr7* while repressing *Gzmb, Itgae* and *Cxcr6*— thus painting KLF2 as a critical director of the LN_CM_ state by enforcing circulation machinery while repressing resident memory transcripts (**Table S1**).

In summation, LN_CM_ are governed by core transcription factors promoting circulation. Both T_RM_ populations lacked these core transcription factors, underscoring the idea that downregulation of the genetic programs facilitating T cell trafficking through blood and lymph is a fundamental hallmark of T_RM_. Interestingly, Lung_RM_ gene networks drive inhibitory receptor expression. LN_RM_ lack both programs which prevents circulation while potentially maintaining effector capacity and low inhibitory receptor expression.

### LN_RM_ have greater proliferative and effector capabilities than Lung_RM_

Our snRNA+ATAC sequencing indicated that LN_RM_ were highly poised for cytotoxicity whereas Lung_RM_ expressed elevated levels of inhibitory receptors, likely due to secondary antigen encounter, suggesting reduced effector capabilities. We questioned whether these epigenetic or transcriptional differences translated to disparities in biophysical or functional properties of memory T cells. We measured the buoyant mass of individual cells from flow sorted memory populations using Suspended Microchannel Resonator (SMR) technology^34^. Buoyant mass measurements were highest for T_CM_(CD69^-^CD62L^+^), lowest for Lung_RM_ (CD69^+^CD103^+^), and intermediate for LN_RM_(CD69^+^CD103^+^) (**Fig 6A**). In prior studies, increased T cell buoyant mass positively correlated with increased proliferative capacity^35^ which is an essential facet of T cell responses to antigen encounter. To directly test proliferative capacity, we labeled flow sorted OT-I memory T cell populations with Cell Trace Violet (CTV) and cultured them *ex vivo* with SIINFEKL(OVA)-pulsed splenocytes for 48 hours (**Fig 6B**). T_CM_ cells were the most proliferative, while Lung_RM_ cells were the least proliferative of the evaluated populations. Consistent with our buoyant mass measurements, LN_RM_ T cells showed an intermediate proliferative capacity (**Fig 6C,D**). These results indicate that the proliferative capacity of resident memory T cells varies by tissue.

**Figure 6.**
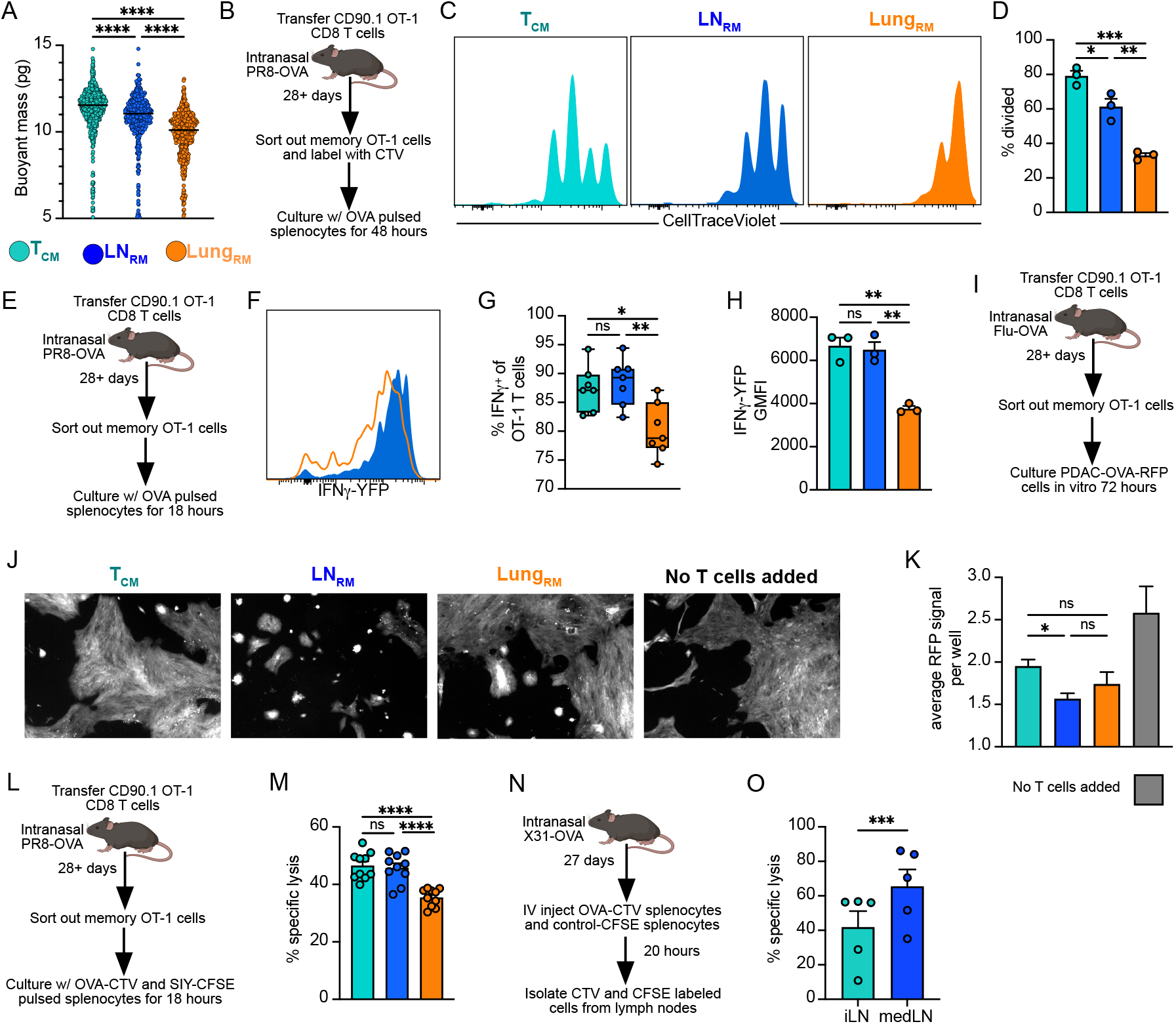
LN_RM_ have greater proliferative and effector capabilities than Lung_RM_. (A) Buoyant mass of sorted memory OT-I T cells 75 days post PR8-OVA. Data are from 1 mouse and representative of 3 mice. T_CM_=CD90.1^+^CD69^-^CD62L^+^ from spleen. LN_RM_=CD90.1^+^CD69^+^CD103^+^ from medLN. Lung_RM_=CD90.1^+^CD69^+^CD103^+^ from lung. (B) Experimental setup for (C-D). (C-D) Representative flow plots and quantification of memory OT-I T cells labeled with CTV and cultured ex vivo with peptide-pulsed splenocytes. Data are representative of two experiments. (E) Experimental setup for (F-H). (F-H) Representative flow plots and quantification of IFNγ production by memory OT-I T cells. Data are representative (F-G) or combined (H) from two experiments. (I) Experimental setup. (J-K) Representative imaging and quantification of fluorescent cancer cells (PDAC: pancreatic ductal adenocarcinoma) cultured with memory T cell populations. Cells were imaged and fluorescence of cancer cells was quantified using an Incucyte S3. Data are combined from 3 separate experiments with n=2-5 per experiment and each n represents one well of T cells and cancer cells. (L) Experimental setup for (M). (M) Quantification of in vitro cytotoxicity assay. Data are combined from 2 experiments. (N)Experimental setup for (O). (O)Quantification of in vivo cytotoxicity assay. Data are representative of 3 experiments with 3-5 mice per experiment.

In response to antigen encounter, memory CD8^+^ T cells exert effector functions via direct killing of target cells and production of cytokines such as IFNγ. Given the importance of cytokine production in secondary immune responses, we measured IFNγ production by memory T cells using an IFNγ-YFP transgene reporter which accurately reports IFNγ protein production^36^. Naïve IFNγ-YFP OT-I T cells were transferred into naïve B6 mice before PR8-OVA infection followed by at least 28 days of rest. Flow sorted IFNγ-YFP OT-I memory T cell populations were restimulated overnight with OVA-pulsed splenocytes (**Fig 6E**). While most T cells were able to produce IFNγ, Lung_RM_ produced significantly less than T_CM_ and LN_RM_ (**Fig 6F-H**). These findings further support the notion that the lack of inhibitory receptor expression in LN_RM_ preserves cytokine production competency compared to Lung_RM_ counterparts.

To test the cumulative effector capabilities of memory T cells we flow sorted memory OT-I T cells and cultured them with a pancreatic ductal adenocarcinoma cell line expressing OVA and red fluorescent protein (PDAC-OVA-RFP) (**Fig 6I**). We measured the total number of PDAC-OVA-RFP cells using an Incucyte S3, by tracking fluorescent signal from PDAC-OVA-RFP over time. As expected, the addition of any memory T cell population reduced the prevalence of PDAC-OVA-RFP, though it was unclear if the reduction was a consequence of cytokine production or a result of direct cell killing. Interestingly, the addition of LN_RM_ resulted in the lowest level of detectable PDAC-OVA-RFP cells demonstrating that LN_RM_ excel at performing classical effector functions of CD8^+^ T cells compared to T_CM_ and Lung_RM_ (**Fig 6J, K**). To more directly assess killing capacity, we utilized an *in vitro* cell killing assay using peptide-loaded splenocytes. Splenocytes from naïve mice were pulsed with OVA or SIY (irrelevant antigen) and labeled with CTV and CFSE respectively, before culturing with flow sorted memory OT-I T cell populations for 18 hours (**Fig 6L**). Surprisingly, T_CM_ and LN_RM_ lysed target cells at similar levels while Lung_RM_ resulted in the lowest level of specific lysis of target cells (**Fig 6M**). We hypothesized that LN_RM_ may contribute to host defense by rapidly killing target cells in the medLN. After an in vivo killing assay in X31-OVA memory mice (**Fig 6N**), OVA loaded cells were more rapidly killed in the medLN than the iLN (**Fig 6O**). As such, LN_RM_ in the medLN may not necessarily kill target cells more efficiently than T_CM_. Because the medLN contains both resident and circulating memory while the iLN contains only circulating memory, the medLN benefits from an additive effector capacity that facilitates direct cell killing of antigen bearing cells. Together, these results reveal that tissue context fundamentally shapes the physical and functional capacity of memory T cells, thereby uncoupling the concept of “residency” from uniform functionality.

## Discussion

Our findings indicate that influenza infection in the respiratory tract seeds functionally divergent resident memory populations in the lung and lung-draining lymph node. Lung_RM_ strikingly resemble exhausted T cells, characterized by quantitative waning in the absence of antigen and expression of multiple inhibitory receptors. More importantly, they exhibit diminished effector capacity, as evidenced by impaired proliferation, cytokine production, and cytotoxicity. Conversely, LN_RM_ are numerically stable and retain effector functions, highlighting a tissue specific divergence in resident memory T cell quality. This variance underscores the need to consider tissue context when designing vaccines and immunotherapies.

The respiratory tract represents one of the largest mucosal barriers and is a common site of infection, raising the critical question of why Lung_RM_ are defined by poor persistence and reduced effector functions. One prevailing teleological explanation is that respiratory functions in the lung have relatively low tolerance for inflammation. However, direct experimental evidence that Lung_RM_ cause aberrant inflammation is lacking. Moreover, this rationale is challenged by T_RM_ in the brain, another tissue with low tolerance for inflammation, where T_RM_ remain numerically stable for over 200 days after LCMV infection^37^. Notably, brain T_RM_ are also high for PD-1 expression^38^, suggesting that tissue-specific adaptations dictate T_RM_ persistence and quality as opposed to a universal requirement to minimize inflammation.

Mechanistically, the transcription factors CREM and SMAD3 appear to be partially responsible for high inhibitory receptor expression and reduced persistence respectively. Both CREM and SMAD3 eRegulons were highly active in Lung_RM_. SCENIC+ analysis inferred CREM as a driver of *Pdcd1, Icos, Ctla4*, and *Havcr2*, which have all been shown to limit CD8^+^ T cell effector function^39^. CREM expression is induced by both TCR signaling and IL-15 sensing and genetic ablation of *Crem* enhances accessibility of *Gzmb*, and *Prf1* loci, further implicating CREM as a negative regulator of lymphocyte effector function^40^. Casp8ap2 (encoding Caspase 8 associated protein 2 or FLASH) was predicted to be upregulated by SMAD3 and is heavily involved in Fas mediated apoptosis. TGF? signaling, which is essential for optimal T_RM_ formation^41^, also prompts activation of SMAD2 and SMAD3 to induce apoptosis of CD8^+^ T cells after listeria infection in mice^42^. Together, these findings support a model in which sustained antigen recognition and cytokine signaling within the infected lung drive CREM and SMAD3 transcriptional programs that limit the persistence and functionality of Lung_RM_.

It was previously observed that LN_RM_ generated after LCMV infection, were predominantly positioned in the subcapsular sinus^7^. Our analyses of the lung-draining lymph node after influenza infection demonstrate a more homogenous distribution of LN_RM_, indicating that the localization of LN_RM_ is not fixed but rather, is highly dependent upon inflammatory conditions (LCMV vs influenza) or precisely which lymph node (inguinal vs mediastinal) the cells are present within. It should be noted that our analyses were restricted to frozen sections of lymph nodes and thus, it is unclear to what extent LN_RM_ migrate between compartments of the lymph node or if they remain stationed within a single compartment. Intravital microscopy is likely best suited to answering these questions but is limited by problems in distinguishing between circulating and resident memory and difficulties in following individual cells over long periods of time in live mice.

While spatial positioning of LN_RM_ appears flexible, their formation, at least in lung-draining lymph nodes, appears highly conserved. Indeed, LN_RM_ formed across multiple antigen specificities, several strains of influenza, and from every precursor population examined. Formation of LN_RM_ in the lung-draining lymph node does not appear to be influenza-specific either, as evidenced by murine experiments with LCMV and vaccinia infections ^9,16,43^. It is difficult to compare the magnitude of LN_RM_ across multiple pathogens, variable amounts of virus and varying number of TCR-tg T cells. With those caveats, influenza appears to seed a surprisingly high number of LN_RM_ with over 70% of influenza-specific T cells expressing canonical T_RM_ markers CD69 and CD103, 21 days post infection. These percentages dwarf those seen in other lymph nodes after vaccinia and LCMV^6,9^. Why influenza seeds such a high degree of LN_RM_ is deserving of further research given the heightened functionality and durability of LN_RM_ compared to Lung_RM_.

Influenza-derived LN_RM_ outcompeted Lung_RM_ in every functional assay we performed. It is unclear if LN_RM_ are universally superior to T_RM_ in non-lymphoid tissues or if this finding is unique to the lung and/or influenza setting. It is possible that prolonged antigen recognition in barrier tissues, relative to the draining lymph node, drives long-term reduction in effector capabilities. In support of this idea, we observed elevated expression of PD-1 and TIM3 on Nur77-GFP^+^ T cells in the lung but found minimal Nur77-GFP expression in the lung-draining lymph node. Alternatively, lymph nodes may foster superior T cell function due to the cytokine milieu or availability of metabolites. These possibilities are not mutually exclusive.

These data support a model of regional immunity defined by compartmentalization of effector capabilities, spatial localization and migration kinetics. While Lung_RM_ provide a rapid front-line defense, their numerical attrition and functional decline potentially represent an evolutionary safeguard to preserve the lung mucosa from immunopathology. In contrast, the draining lymph node acts as a sanctuary by fostering numerically stable and functionally competent memory T cells outside the hostile environment of the lung. This dichotomy suggests that long-term mucosal immunity does not solely rely on the indefinite persistence of resident memory cells in the lung, but on a tiered system with the draining lymph node safeguarding a more functional resident memory pool required for optimal adaptive immune responses upon reinfection.

## Acknowledgements

We thank: Ryan Langlois and Susan Kaech for providing PR8 and X31 viruses respectively; S. Levine at the MIT BioMicro Center for sequencing support; Emma Dawson for providing the PDAC-OVA-RFP cell line; Tyler Jacks for Incucyte equipment; Melissa Duquette for help with mouse colony management. Figures were made, in part, using Biorender.

S.S. is supported by American Cancer Society ACS RSG-23-863228-01-IBCD and NCI 1R01CA273819-01A1. T.A.H. is a Ludwig Center at MIT’s Koch Institute postdoctoral fellow and member of the Convergence Scholars Program at MIT. J.M.S. is a Cancer Prevention and Research Institute of Texas (CPRIT) Scholar in Cancer Research. J.M.S. is an Andrew Sabin Family Foundation Fellow at The University of Texas MD Anderson Cancer Center. S.M. is supported by Virginia and D.K. Ludwig Fund for Cancer Research and MIT Center for Precision Cancer Medicine. J.C.L. is supported by a SPARC grant from the Broad Institute and the Daniel I.C. Wang (1959) Faculty Research Innovation Fund at MIT. Z.J.R is supported by American Cancer Society postdoctoral fellowship (PF-24-1244739-01-IBCD).

## Ethics Declarations

S.S. is a scientific advisory board member for Related Sciences, Arcus Biosciences, Ankyra Therapeutics, Arpelos Biosciences, and Repertoire Immune Medicines. S.S. is a co-founder of Danger Bio and is a consultant for TAKEDA and Merck. S.S. receives funding for unrelated projects from Merck. S.M. is a founder of Travera and Affinity Biosensors. J.C.L. has interests in Amplifyer Bio, Sunflower Therapeutics PBC, Honeycomb Biotechnologies, OneCyte Biotechnologies, QuantumCyte, and Repligen.

## Author contributions

T.A.H. and S.S. conceived the study and wrote the original manuscript. Experiments were conducted by: T.A.H., S.D., M.Y.C., V.B., F.C., and T.B. Sequencing analysis was conducted by T.A.H. and Z.R. Experiments were designed by: T.A.H., J.M.S and S.S. Funding was acquired by S.S. All authors read and contributed to editing the manuscript.

## Methods

### Mouse strains

Female C57BL/6J mice from 6-8 weeks of age were purchased from Jackson Laboratory and maintained in specific-pathogen free rooms. Thy1.1 (strain: 000406), OT-I (strain:003831), Nur77-GFP (strain: 016617), IFNγ-YFP (strain: 017581) mice were purchased from Jackson Laboratory and bred in house.

### Longitudinal antibody labeling

Longitudinal antibody labeling was performed in one of two ways. For Fig1B-E, mice received 3ug of anti-CD45.2-PE via retro-orbital injection 24 and 18 hours before tissue harvest. For Fig1F-H, mice received 5ug of anti-CD45.2-PE via retro-orbital injection and 5ug of anti-CD45.2-AF657 via intraperitoneal injection 24 hours before tissue harvest.

### Tissue processing, flow cytometry and cell sorting

Lymph nodes were smashed along scored wells of 12 well plates and digested in 1mg/mL of Collagenase D with 50U/mL deoxyribonuclease in Hank’s balanced salt solution. Lungs were removed from mice and placed in 5mL of the same digestion media in a gentleMACS C tube. Lungs were mechanically digested using a gentleMACS Octo Dissociator with Heaters. Samples were first digested with the m_impTumor_01_01 program, followed by the 37C_Multi_E_01 program. Spleens were smashed over a 70µm filter. Red blood cells in spleens and lungs were lysed using Ammonium-Chloride-Potassium (ACK) lysis buffer. Single cell suspensions were stained with CD8a (53-6.7), CD44(IM7), CD90.1(HIS51 or OX-7), CD103(2E7 or M290), CD62L(MEL-14), CD45(30-F11), PD-1(RMP1-30 or 29F.1A12), GranzymeB (GB11), CD69(H1.2F3), TIM3 (RMT3-23), in PBS with 1% bovine serum albumin (BSA). Dead cells were identified using eBioscience Fixable Viability Dye eFluor 780 and stained samples were run on a BD Symphony A3. A FACS Aria II with a 70um nozzle was used for cell sorting.

### Human thoracic lymph node analysis

Data from the Human Cell Atlas at (https://cellxgene.cziscience.com/collections/cc431242-35ea-41e1-a100-41e0dec2665b) was used to calculate the prevalence of CD8^+^ T cell subsets in human thoracic lymph nodes. Analysis was restricted to the T cell subset dataset and consisted of single cell RNA and surface protein expression. Naïve, T_CM_, T_EM_ and T_RM_ status were distinguished by gene expression and detection of surface protein. Definitions were based upon MultiModal Classifier Hierarchy (Wells et al 2025) and the authors definitions. CD8^+^ MAIT cells were excluded from analysis. Briefly, T_RM_ were defined by CD69 surface protein detection, or at least two of the following: *Itgae, Itga1, Cxcr6*, CD103 or CD49a surface protein detection. Naïve CD8^+^ T cells were defined by CD62L and CD45RA surface protein detection. T_CM_ were defined by CD62L, and one of CD45RO, CD95 or *Ccl5*. T_EM_ were defined by low expression of CD62L and CD45RA. T_EMRA_ were defined by CD45RA and 1 of (*Ccl5, Fcgr3a, Gzmb*, G*zmk*, G*zma, Ifng, Pdcd1*, CD57, *Fgfpb2*, or KLRG1). Subset definitions are discussed further in Wells et al. 2025.

### Immunofluorescent imaging

Mediastinal lymph nodes were placed in optimal cutting temperature (OCT) compound. Samples were frozen on an isopentane bath over dry ice. After freezing, 7-10 µm sections were cut and fixed in ice cold acetone for 10-15 minutes. Sections were blocked in PBS with 1% BSA for 10 minutes. Endogenous biotin and avidin were blocked using ReadyProbes Avidin/Biotin Blocking Solution (cat: R37627) if biotinylated antibodies were used. Staining was performed for 3-4 hours at room temp in the dark. Staining solution was removed, and #1.5 coverslips were mounted using Prolong Diamond mounting media. Images were acquired using a TissueFAXS SL fluorescent slide scanner. OT-I T cells expressing CD90.1 in the lymph node were manually counted in FIJI. Cells were placed in one of the four following categories (1) Subcapsular sinus (SCS), (2) in direct contact with LYVE1^+^ vessels outside the SCS, (3) in follicles or (4) within the T cell zone or interfollicular areas. Area of regions was calculated in FIJI.

### Buoyant mass measurements

The SMR is a cantilever-based system with an internal microchannel. The cantilever vibrates at a baseline resonant frequency and as a cell flows through the channel, the frequency changes. The detected frequency change is correlated with the buoyant mass of the cell. Further details regarding operation and fabrication can be found in Burg et al^34^. Regulators, valves, and data acquisition are controlled by custom software coded in LabVIEW 2020 (National Instruments).

At 75 days post PR8-OVA infection, flow sorted memory OT-I T cells were isolated as previously described. Before measurement collection, the SMR was cleaned with 0.05% Trypsin-EDTA (Invitrogen, 25300054) for 10 minutes, then 10% bleach for 3 minutes, and finally DI-H_2_O for 3 minutes to remove any cell debris. Measurements were obtained in Flow Cytometry (FACS) Staining Buffer (Rockland, MB-086-0500) at room temperature for about 30 minutes per sample and channels were flushed with buffer in between samples.

### In vitro activation and cytotoxicity assays

For in vitro proliferation assays, memory CD90.1 OT-I T cells were sorted out of immune chimeras and labeled with Cell Trace Violet (Invitrogen C34571) as per manufacturer recommendations and then cultured in vitro with SIINFEKL(OVA) pulsed splenocytes from naïve mice at 37C.

For in vitro cell killing assays, splenocytes were pulsed with either SIINFEKL or SIYRYYGL and then labeled with either CTV or CFSE and then mixed at a 1:1 ratio. Splenocytes were pulsed with 1-2uM peptide for 1-3 hours at 37C. 3000-10000 memory OT-I T cells were plated per well with OVA or SIYRYYGL(SIY) pulsed splenocytes in a 96 well plate at a ratio of 1:3 (OT-I: OVA-pulsed splenocyte).

For in vivo killing assays, splenocytes were similarly pulsed with OVA or no peptide control after CTV and CFSE labeling. Pulsed and labeled splenocytes from CD45.1 mice were injected into X31-OVA mice at 27-28 days post infection. Splenocytes were also transferred into naïve mice for a control. 20-22 hours later, inguinal and mediastinal lymph nodes were isolated. Specific lysis of target cells was calculated relative to lymph nodes from control mice. To investigate T cell killing of cancer cells, 2000-4000 memory OT-I T cells were plated in a flat bottom 96 well plate with twice as many PDAC-OVA-RFP cells. Fluorescence of PDAC-OVA-RFP cells was captured over time with an Incucyte S3.

### Single nulcei RNA+ATAC sequencing

Live, extravascular CD44^+^CD8^+^CD90.1^+^ memory OT-I T cells were sorted out of lungs or medLNs from 18 immune chimeras 46 days post PR8-OVA infection. Cells from the same tissue were pooled together. Nuclei isolation was adapted from 10x protocol CG000365 and RNAse inhibitor (10x) was added to prevent RNA degradation. Single nuclei suspensions of cells from lungs or medLNs were submitted to the MIT BioMicro Center for library preparation and sequencing. Isolated nuclei were counted on a Luna-fl dual fluorescence cell counter. Single cell coupled ATACseq and nuclearRNAseq were generated following 10X Genomics standard multiome kit methods using 7 cycles of PCR for ATACseq and cDNA generations. Final libraries were generated using modified oligonucleotides to include Singular S1/S2 anchors in places of the Illumina P5/P7 sequences. snATACseq and snRNAseq libraries were each sequenced on a full SingularG4 F2 flowcell. Data were analyzed in R studio using Seurat and Signac packages. Low quality nuclei data was removed under the following criteria: (nCount_ATAC < 100000), (nCount_RNA < 25000), (nCount_ATAC > 1800), (nCount_RNA > 1000), (nucleosome_signal < 2), (TSS.enrichment > 1), (percent.mt < 25). RNA sequencing from the medLNs and lungs were merged using the Seurat merge function. A resolution of 0.2 was used for clustering and a small (<200) number of cells expressing B cell receptor transcripts was removed from subsequent analysis. The chromVAR function from Signac was used to determine motif activity. Fold changes >1 with an adjusted P value of <0.05 were considered differentially accessible regions or differentially expressed genes.

Enhancer-driven gene regulatory network activity was determined using SCENIC+^33^. SCENIC+ was run as described at https://scenicplus.readthedocs.io/en/latest/index.html using default parameters. The pipeline was run on 6,686 paired snATACseq/RNAseq cells. For snATACseq, the mm39 genome was used for chromosome sizes and genome annotation. Topic modeling was performed using Parallel LDA and 25 topics were selected based on model convergence. For snRNAseq, gene expression matrices were processed in Seurat as previously described. The murine v10 cisTarget database was used. The pipeline identified 25 direct regulons and 37 extended regulons. Regulon graphs were constructed based on TF-gene importance scores (importance_TF2G, outer radii) and TF-region correlation scores (importance_x_rho, inner radii).

## Data and Materials Availability

Raw data of snRNA+ATAC sequencing has been deposited at National Center for Biotechnology Information (www.ncbi.nlm.nih.gov) and is available upon request.

## Statistical analysis

Ideal sample sizes for experiments were based on prior experience. Statistical significance was calculated using GraphPad Prism version 10 or standard Seurat/Signac functions in R studio. Paired or un-paired Student’s t tests were used for comparisons between two groups. One-way analysis of variance was used to compare more than two groups*P ≤ 0.05, **P ≤ 0.01, ***P ≤ 0.001, ****P ≤ 0.0001. P values greater than 0.05 were considered not significant (NS). Error bars show the mean + SEM.

