## Supplemental Figures for "Lymph node resident memory T cells retain effector capabilities by evading lung resident memory dysfunction"

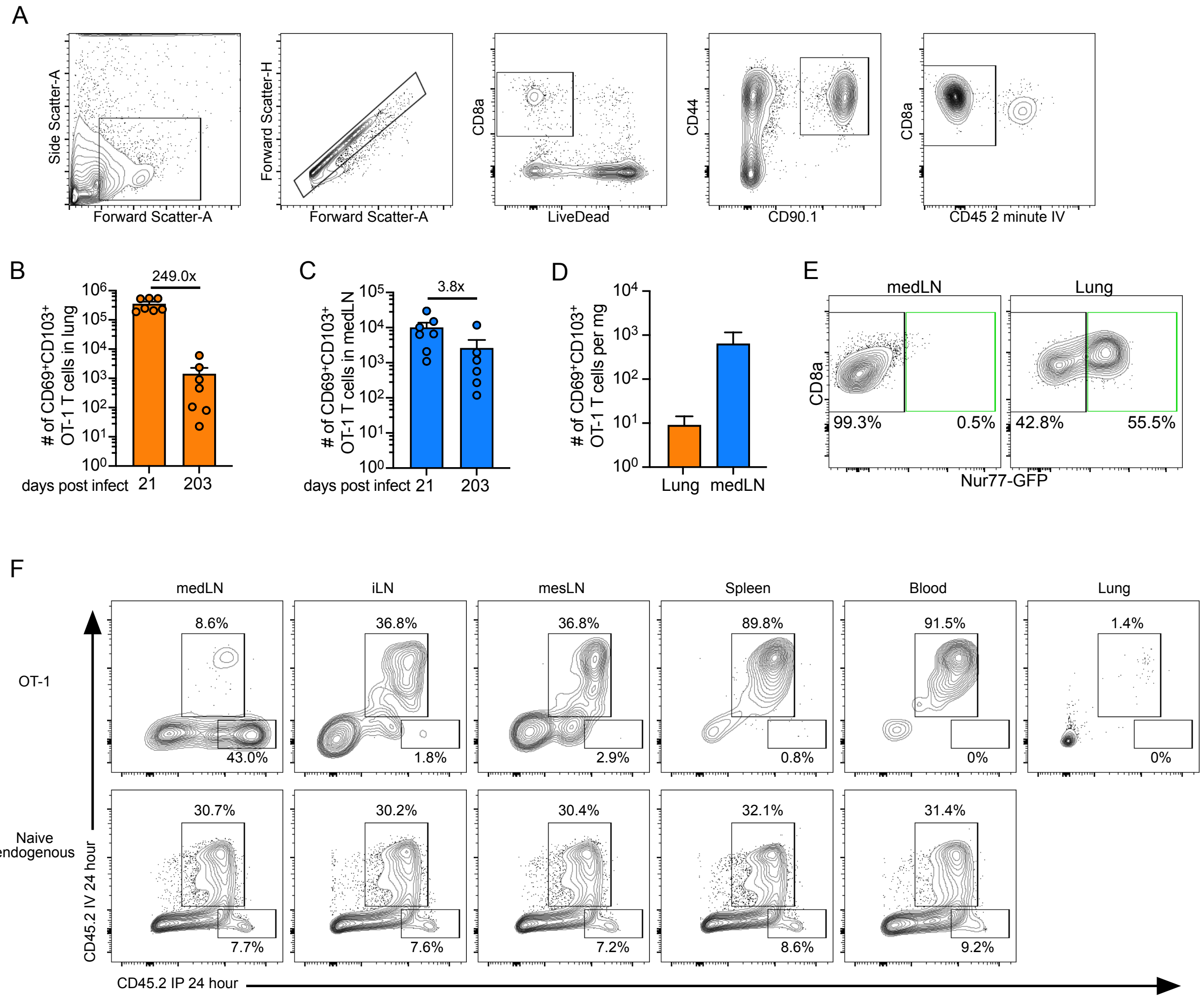


### Supplementary Figure 1

(A) Representative gating scheme of lung 21 days post intranasal PR8-OVA infection. (B-C) Quantification of $CD69^{+}CD103^{+}$ OT-1 T cells. (D) Quantification at 203-326 days post infection. Data are combined from at least two experiments with n=2-4 per experiment (E) Representative flow plots related to Fig 1C. (F) Representative flow plots related to Fig 1F-G. Gated on $CD8^{+}$ $CD90.1^{+}$ OT-I T cells (top) or endogenous $CD8^{+}CD90.1^{-}CD44^{-}$ (bottom). All plots are also gated on extravascular (negative for 2-minute CD45 IV) cells except for the blood.

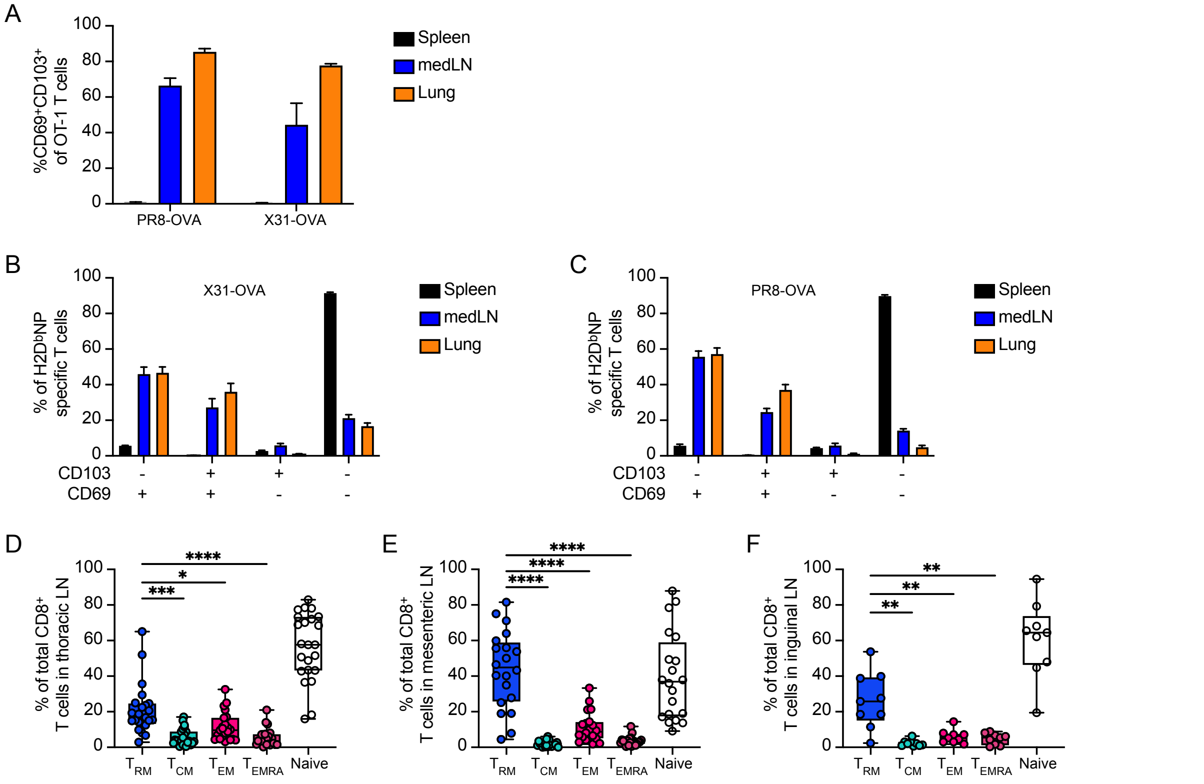


**Supplementary Figure 2**

Related to figure 2. (A-C) Phenotype of extravascular OT-I or tetramer$^+$ T cells 26-27 days post PR8-OVA or X31-OVA. N=3-5 and data are representative of two experiments. Analysis of scRNA+CITEseq of human lymph nodes adapted from Wells et al. 2025. See methods for subset definitions.

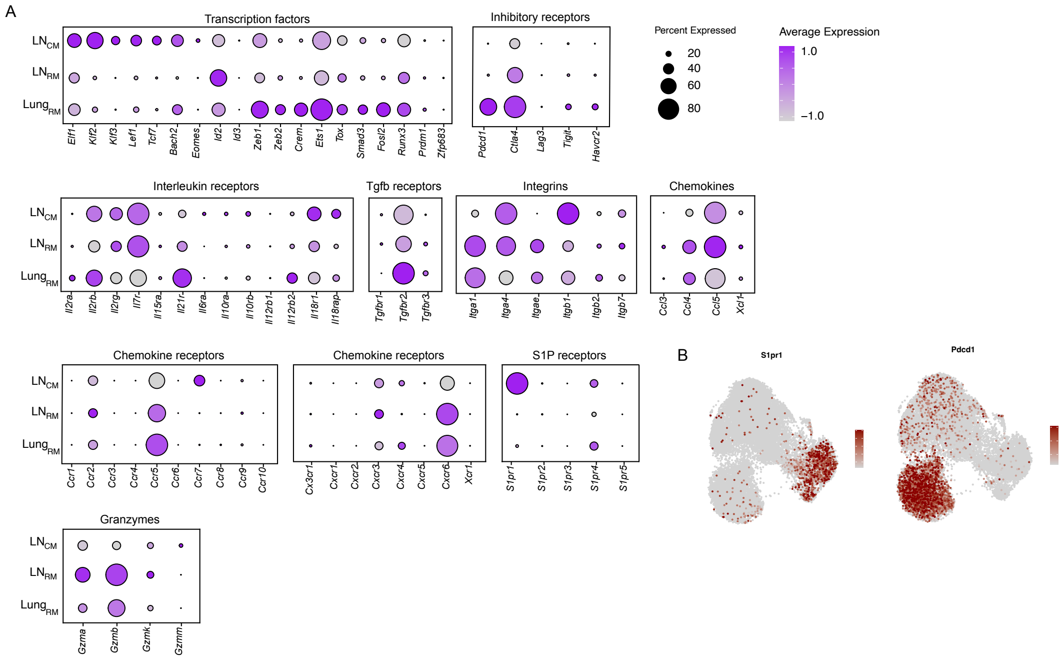


### Supplementary Figure 3

Related to figure 4. (A) Dotplots showing scaled average expression of indicated genes. (B) Feature plot of *S1pr1* transcription in UMAP space.

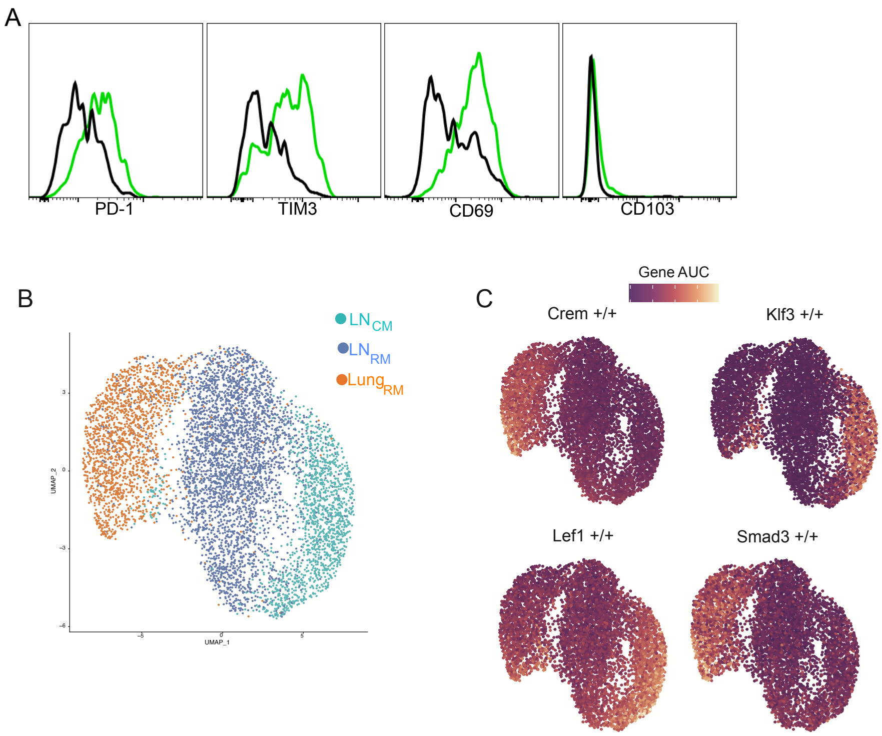


**Supplementary Figure 4**

Related to figure 5. (A)Expression of indicated markers on Nur77-GFP$^-$ (black) or Nur77-GFP$^+$ (green) OT-I T cells from the lung 7 days post intranasal PR8-OVA infection. (C) SCENIC+ analysis was run on cells from figure 4 dataset. Cells were clustered based on eRegulon activity and plotted in UMAP space. (D) eRegulon activity of indicated transcription factors.
